# Deconvolving multiplexed protease signatures with substrate reduction and activity clustering

**DOI:** 10.1101/564906

**Authors:** Qinwei Zhuang, Brandon Alexander Holt, Gabriel A. Kwong, Peng Qiu

## Abstract

Proteases are pleiotropic, promiscuous enzymes that degrade proteins and peptides, which drive important processes in health and disease. The ability to quantify the activity of protease signatures by sampling with Massively Multiplexed Activity (MMA) libraries will provide unparalleled biological information. Under such a framework, a designed library of peptide substrates is exposed to a cocktail of proteases, the cleavage velocity of each substrate is measured, and individual protease activity levels are inferred from the data. Previous studies have developed individual protease sensors, but multiplexed substrate cleavage data becomes difficult to interpret as the number of cross-cutting proteases increases. Computational methods for parsing this data to estimate individual protease activities primarily use an extensive compendium of all possible protease-substrate combinations, which require impractical amounts of training data when scaling up to MMA libraries. Here we provide a computational method for estimating protease activities efficiently by reducing the number of substrates and clustering proteases with similar cleavage activities into families. This method is scalable and will enable the future use of MMA libraries with applications spanning therapeutic and diagnostic biotechnology.

## 1 Introduction

Proteases are multifunctional enzymes that cleave peptide bonds and are responsible for maintaining health in processes ranging from immunity to blood homeostasis, but are also drivers of diseases, including cancer and sepsis (*1-10*). Certain aspects of protease biology remain poorly understood, such as the mechanisms regulating substrate specificity *in vitro* vs. *in vivo* (*11*). Libraries of protease sensors are required to perform these experiments, which led to the development of more specific and efficient probes. There are ∼550 known proteases expressed by the human genome, which requires a substrate library of approximate magnitude to completely resolve an individual’s protease landscape, assuming one protease cuts one substrate. Next Generation Sequencing technologies provide the ability to rapidly assess protein *abundance* on this scale, but previous studies show a lack of correlation between expression and enzymatic activity (*12-14*). The quantification of protease *activity* on this scale in humans will provide copious amounts of unseen biological information, leading to improved diagnostic and therapeutic technologies.

For this reason, countless platforms have been developed to sense and modulate protease activity *in vivo* and *in vitro*, with the potential to extract useful physiological information (*10, 15-30*). These technologies require experimental knowledge of protease-substrate specificity (*11*), which is difficult to completely measure because proteases are promiscuous (*8*), enabling them to cut multiple different substrate sequences. Therefore, independent protease signatures become convolved when multiplexing a library of sensors, making it difficult to quantify the relative activity of each protease (*31*). Previous studies have successfully developed computational algorithms to parse these signatures (*32*), but these methods may become complicated when applied to proteases with similar substrate signatures.

Here, we present a method for deconvolving protease signatures, which requires limited prior knowledge of protease-substrate specificity. To overcome the challenge of scaling to all physiological proteases, we use this method to improve experimental design by reducing the size of the substrate library. Furthermore, we cluster proteases with similar substrate activities into families, while maintaining high estimation accuracy. Under this framework, we lay the groundwork for understanding multiplexed protease-substrate signatures on a large scale, enabling the use of Massively Multiplexed Activity (MMA) libraries.

## 2 Method

To improve experimental design for deconvolving protease composition of protease mixtures, we developed pipelines for estimating kinetic parameters from real experimental data, and simulating *in silico* experimental data. In this Method section, we firstly introduce individual components of the pipeline (Method 2.2-2.5). We then apply the pipeline to optimize the selection of substrates and cluster proteases into families (Method 2.6).

The overall strategy for the deconvolution analysis consists of two optimization steps. The first step learns the cleaving dynamics of every combination of one protease and one substrate by optimizing kinetic parameters for a modified Michaelis-Menten model (*33, 34*) (see details in Method 2.2.2 and 2.4.1). The second step applies the kinetic parameters learned in the first step to estimate the mixing coefficients, representing the concentrations of proteases in the protease mixtures to be deconvolved (see details in Method 2.4.2). In the case where a sufficient number of substrates are measured to deconvolve all individual proteases in a mixture, we screen for the optimal subset of substrates in order to reduce the required number of substrates. When highly correlated proteases exist in the mixture, which would require an impractically large number of substrates for deconvolution, we cluster the proteases into families via hierarchical clustering to enable deconvolution based on a reasonable number of substrates and achieve a higher accuracy at a lower resolution (Method 2.6).

### 2.1 Recombinant Protease Activity Assay

To obtain the recombinant protease activity data, we first conjugated seven different c-terminus cysteine synthetic peptide substrates to amine functionalized 2 μm magnetic microparticles with SIA, an amine-thiol crosslinker. The n-terminus of the peptides each contains one of seven unique glu-fib mass barcodes (**Table 1**). We then incubated a cocktail of these seven substrates with each of the seven recombinant proteases individually at 37°C on a spinner. At various time points between 0 and 400 minutes, we used a magnetic separator to remove the microparticles from the supernatant, which contains the cut substrates plus mass barcodes. Mass spectrometry was performed to quantify the amount of cut substrate at each time point.

**Table 1.**

| Substrate Name | Peptide sequence (N terminus on left) | Modifications |
| --- | --- | --- |
| CC01 | e(*aa)(*aa)ndneeGFFsAr(ANP)K(5-FAM)GGLQRIYKC | 1st *aa= Gly(13C2); 2nd *aa=Val(U13C5,15N) |
| CC02 | eG(*aa)ndneeGF(*aa)s(*aa)r(ANP)K(5-FAM)GGKSVARTLLVKC | 1st *aa= Val(U13C5,15N); 2nd *aa=Phe(15N); 3rd<br>*aa=Ala(15N) |
| CC03 | e(*aa)(*aa)ndneeGFFs(*aa)r(ANP)K(5-FAM)GGQRQRIIGGC | 1st *aa= Gly(U13C2,15N); 2nd *aa=Val(15N); 3rd<br>*aa=Ala (U13C3,15N) |
| CC04 | e(*aa)Vndnee(*aa)FFs(*aa)r(ANP)K(5-FAM)GGKYLGRSYKVC | 1st *aa= Gly(13C2); 2nd *aa=Gly(13C2); 3rd<br>*aa=Ala(U13C3,15N) |
| CC05 | eGVndnee(*aa)(*aa)Fs(*aa)r(ANP)K(5-FAM)GGGLQRALEIC | 1st *aa=Gly(U13C2,15N); 2nd *aa=Phe(15N); 3rd<br>*aa=Ala(U13C3,15N) |
| CC06 | e(*aa)(*aa)ndnee(*aa)(*aa)(*aa)s(*aa)r(ANP)K(5-FAM)GGKTTGGRIYGGC | 1st *aa=Gly(13C2); 2nd *aa=Val(U13C5,15N); 3rd<br>*aa=Gly(U13C2,15N); 4th *aa=Phe(15N); 5th<br>*aa=Phe(15N); 6th *aa=Ala(15N); still include ANP and<br>K5-FAM |
| CC07 | eG(*aa)ndnee(*aa)(*aa)Fs(*aa)r(ANP)K(5-FAM)GGQARGGSC | 1st *aa=Val(U13C5,15N); 2<sup>nd</sup> *aa=Gly(U13C2,15N);<br>3rd *aa=Phe(15N); 4th *aa=Ala(U13C3,15N) |

### 2.2 Cleaving dynamics approximation model

#### 2.2.1 Michaelis-Menten kinetics as the approximation of cleaving dynamics

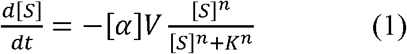

[*S*]: the remaining amount uncleaved substrate, [*S*] ∈ [0,1], [*S*]^*t=0*^= 1

*V*: the maximal rate of the reaction at the saturating substrate concentration

*K*: the substrate concentration when the reaction rate reaches half of *V*

*n*: the order of reaction

[*α*]: concentration of a protease

We use the Michaelis-Menten kinetics as the base model to approximate the cleaving dynamics (*33, 34*). Worth noticing, the mixing coefficient [*α*], the concentration of a protease, is added to the original Michaelis-Menten model. On the left-hand side (LHS) of equation (1), *d[S]/dt* represents the rate of change of the remaining uncleaved substrate. At t=0, *d[S]/dt* is negative, which means that the substrate is being cleaved and will decrease. As [*S*] decreases, the cleaving process slows down until *[S]* arrives at 0, where the reaction stops due to the depletion of the uncleaved substrates. However, in real experimental data, we noticed persistent non-zero saturation levels of uncleaved substrates, which motivated a modification of the model by adding a saturation term *β* (see details in Method 2.2.2).

#### 2.2.2 Modified Michaelis-Menten kinetics with saturation

Equation (2) is the modified Michaelis-Menten model for approximating the changing rate of one uncleaved substrate species when reacting with one protease. *β* is the saturation term representing the concentration of uncleaved substrate at which the reaction stops (when *S* = *β*, the RHS becomes 0). In subsequent discussions, we refer to V, K, n, and *β* as kinetic parameters, and [*α*] as the concentration mixing coefficient.

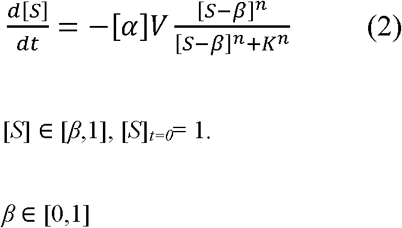

To generalize Equation (2) for modeling the dynamics of substrates cleavage by protease mixtures, subscripts *i* and *j* are introduced in Equation (3), which models the changing rate of the uncleaved substrates when reacting with multiple proteases.

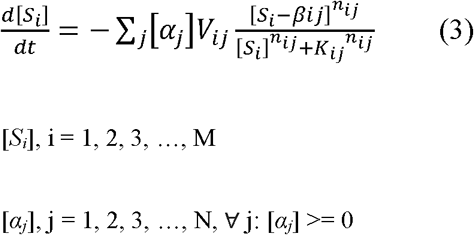

[*S*_*i*_] is the amount of uncleaved substrates of the *i*^*th*^ substrate in the substrate library. [*α*_*j*_] is the concentration of *j*^*th*^ protease in the mixture. Let *M* be the number of substrates and *N* be the number of proteases. Equation (3) assumes that no synergistic or antagonistic effect is involved among various proteases within the protease mixture when cleaving substrates.

### 2.3 Simulating *in silico* experiments

#### 2.3.1 Simulating data in the single-protease-single-substrate setting

Given [*S*]_*t=0*_= 1, [*α*] = 1, and a set of specific kinetic parameter values (*V, K, n, β*), the amount of remaining uncleaved substrate [*S*]_*t=t*_*z*__ at a specific time *t = t*_*z*_ can be calculated by numerically solving equation (2). The kinetic parameter values are either randomly generated or estimated from real experimental data under single-protease-single-substrate setting (Method 2.4.1). Let *Q* be the number of measurement time points, *t = t_z_* (*z* = 1, 2,…, *Q*). This simulation generates a *Q* × 1 data vector representing the simulated amounts of the uncleaved substrate at *Q* time points for the single-protease-single-substrate scenario.

#### 2.3.2 Simulating data in the multi-proteases setting

Given a *M*-substrates library and a *N*-proteases mixture, coefficient [*α*_*j*_] as the concentration of *j*^*th*^ protease in the mixture, and (*V*_*ij*_, *K*_*ij*_, *n*_*ij*_, *β*_*ij*_) as kinetic parameters for the reaction between the *i*^*th*^ substrate and the *j*^*th*^ protease, the amount of remaining uncleaved *i*^*th*^ substrate [*S*_*i*_]_*t= t*__*z*_ at a specific time *t = t*_*z*_ can be calculated by numerically integrating equation (3). The values of the kinetic parameters and mixing coefficients are either randomly generated or estimated from real experimental data (Method 2.4.2). This simulation generates an *M* × *Q* matrix as the simulated data.

### 2.4 Estimating kinetic parameters and mixing coefficients

#### 2.4.1 Estimating kinetic parameters in single-protease-single-substrate setting

For each single-protease-single-substrate combination, we measure/simulate reaction products at *Q* time points after reaction starts, resulting in a data vector ***Y*** with dimension *Q* × 1. In addition to simulated *in silico* experiments (Method 2.3.1), ***Y*** can also be collected from real experiments under the single-protease-single-substrate setting. The problem of estimating the kinetic parameters can be formulated as the following optimization problem:

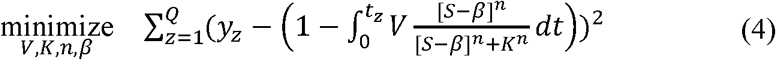

Using the “active-set” algorithm (*35-37*), this optimization problem leads to one set of kinetic parameters that can best fit the data ***Y*** of the specific protease-substrate combination. For a collection of single-protease-single-substrate settings of *M* substrates and *N* proteases, optimization is performed for each of the *M* × *N* combinations, resulting in kinetic parameter matrices (***V***, ***K***, ***n***, ***β***), each of which has a dimension of *M* × *N*.

#### 2.4.2 Estimating mixing coefficients in multi-proteases setting

Once kinetic parameters for all single-protease-single-substrate combinations have been estimated, we move to estimate the mixing coefficients of proteases in a protease mixture. Let *M* be the number of substrates, *N* be the number of proteases in the mixture, and *Q* be the number of measurement times for the reaction between each substrate and the mixture. The problem of estimating the mixing coefficients can be formulated into an optimization problem as follows:

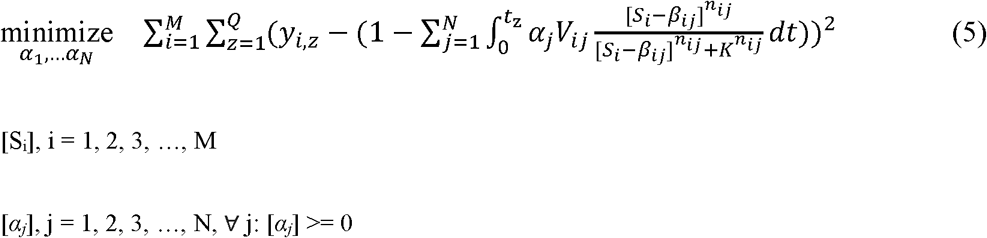

***Y*** is an *M* × *Q* data matrix either generated from *in silico* simulation (Method 2.3.2) or collected from real experiment under the multi-proteases setting. This optimization problem, solved by “active-set” algorithm (*35-37*), generates estimations of the mixing coefficients of proteases in the mixture.

### 2.5 Quantifying estimation accuracy via Root-Mean-Square Error (RMSE)

Once estimated mixing coefficients of a protease mixture have been obtained, the estimation accuracy is evaluated by the root-mean-square error (RMSE) metric. This metric is commonly used in machine learning to quantify accuracy for regression analysis (*38*). An example of quantifying estimation error using RMSE is in **Table 2**.

**Table 2.** The first row is true *α* in P mixtures, of which each has N proteases. The second row is estimated *α*. The RMSEs for individual proteases (R_1_,…, R_N_) are calculated in the third row, and the overall RMSE will be the average of all individual RMSEs. In the simulation setting, P is the number of repetitions we applied. The repetition time is P = 200.

|  | Protease 1 | Protease 2 | ... | Protease N |
| --- | --- | --- | --- | --- |
| True $\alpha$ | $\begin{bmatrix} \alpha_{11} \\ \alpha_{12} \\ \dots \\ \alpha_{1P} \end{bmatrix}$ | $\begin{bmatrix} \alpha_{21} \\ \alpha_{22} \\ \dots \\ \alpha_{2P} \end{bmatrix}$ | ... | $\begin{bmatrix} \alpha_{N1} \\ \alpha_{N2} \\ \dots \\ \alpha_{NP} \end{bmatrix}$ |
| Estimated $\hat{\alpha}$ | $\begin{bmatrix} \hat{\alpha}_{11} \\ \hat{\alpha}_{12} \\ \dots \\ \hat{\alpha}_{1P} \end{bmatrix}$ | $\begin{bmatrix} \hat{\alpha}_{21} \\ \hat{\alpha}_{22} \\ \dots \\ \hat{\alpha}_{2P} \end{bmatrix}$ | ... | $\begin{bmatrix} \hat{\alpha}_{N1} \\ \hat{\alpha}_{N2} \\ \dots \\ \hat{\alpha}_{NP} \end{bmatrix}$ |
| RMSE (Protease) | $R_1 = \sqrt{\frac{\sum_{k=1}^P (\hat{\alpha}_{1k} - \alpha_{1k})^2}{P}}$ | $R_2 = \sqrt{\frac{\sum_{k=1}^P (\hat{\alpha}_{2k} - \alpha_{2k})^2}{P}}$ | ... | $R_N = \sqrt{\frac{\sum_{k=1}^P (\hat{\alpha}_{Nk} - \alpha_{Nk})^2}{P}}$ |
| RMSE (overall) | $R = \frac{\sum_{j=1}^N \sqrt{\frac{\sum_{k=1}^P (\hat{\alpha}_{jk} - \alpha_{jk})^2}{P}}}{N}$ or $R = \frac{\sum_{j=1}^N R_j}{N}$ | | | |

### 2.6 Evaluating deconvolution performance and optimizing substrate selection

Given the kinetic parameter values of the reactions between a set of proteases and a set of substrates, we would like to evaluate whether we can accurately deconvolve mixtures of the proteases by measuring their cleaving activities against the substrates. We first simulate the *in silico* experimental data corresponding to the single-protease-single-substrate scenario (Equation 2) and simulate the *in silico* experimental data for protease mixtures reacting with multiple substrates (Equation 3). We then estimate the kinetic parameter values based on the simulated single-protease-single-substrate data (Equation 4). Finally, we estimate the mixing coefficients based on the estimated kinetic parameters and the simulated experimental data for protease mixtures reacting with multiple substrates (Equation 5) and evaluate the deconvolution accuracy using RMSE. In this analysis pipeline, we choose to estimate the single-protease-single-substrate kinetic parameters because the true kinetic parameter values are often unavailable in practice. Using this pipeline, we can evaluate the expected deconvolution performance for a given set of proteases using a given set of substrates and then derive optimal experimental designs for choosing the most suitable substrates for deconvolving the protease mixtures.

## 3. Results

### 3.1 Recombinant Protease Substrate Specificity

To obtain kinetic protease activity data we incubated 7 serum proteases from the complement and coagulation cascades with 7 protease substrates (**Figure 1**). Proteases displayed different preferences for substrates but were not necessarily linearly independent. Interestingly, proteases from different physiological families showed similar substrate specificity (i.e., MASP2 and CFI). This suggested that computationally-derived families of proteases may not necessarily reflect the biological families. Additionally, each protease showed unique early saturation levels, which is characterized by the parameter *β*.

**Figure 1.**
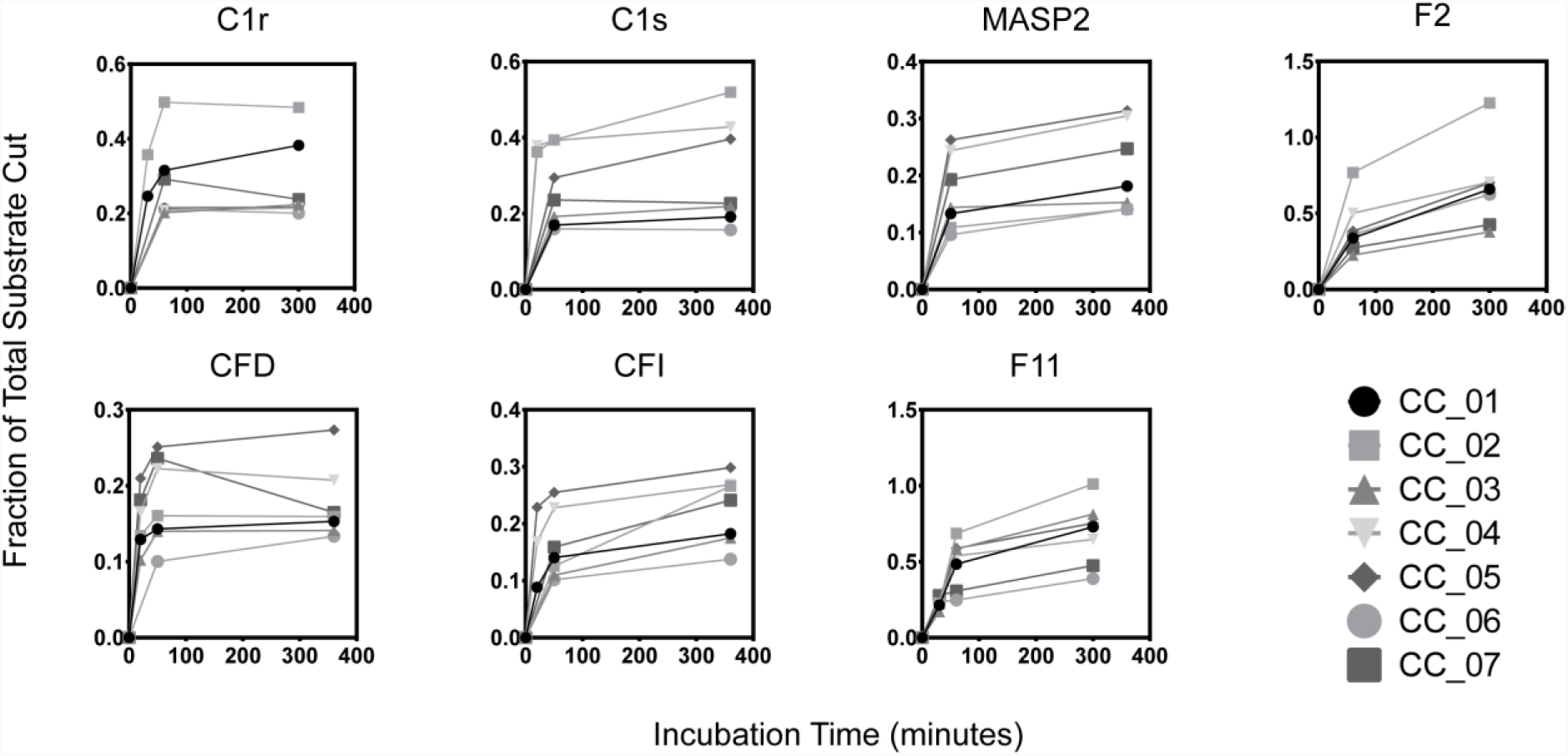
Recombinant protease cleavage assays of seven complement and coagulation cascade proteins. From left to right, top to bottom, abbreviations are: Complement proteins C1r and C1s, MASP2, Coagulation Factor IIa, Complement Factor D, Complement Factor I, and Coagulation Factor XIa. Each trace represents a different peptide substrate (CC01—07).

### 3.2 Validating the RMSE for evaluating protease deconvolution

In order to validate the robustness of RMSE in reflecting estimation accuracy, we simulated a series of 2-protease mixtures with increasing levels of similarity between the two proteases, which represented deconvolution problems with an increasing level of difficulty. In the simulations, the number of observed time points Q was 2, which matched our real experimental setting shown in **Figure 1**. More specifically, we first simulated two proteases (*p*_1_, *p*_2_) by randomly generating their kinetic parameters against multiple substrates. Since the kinetic parameters were randomly generated, these two proteases are independent of each other. We then generated a series of intermediate proteases by linearly combining the two sets of kinetic parameters: *p*_3_ = *λp*_1_ + (1 − *λ*)*p*_2_, *λ* = 0, 0.05, 0.1, 0.15,…,1. After that, we simulated substrate cleavage data of protein mixtures of *p*_1_ and *p*_3_ defined by varying values for *λ*, performed optimizations to estimate the mixing coefficients of the mixtures, and applied RMSE to evaluate the estimation accuracy. Intuitively, the estimation problem is more difficult for cases where the mixed proteases are highly correlated (*λ* close to 1). In addition, we simulated cases with anywhere between 2 to 7 substrates. Generally, the more substrates are measured, the easier it is to deconvolve the protease mixtures. In these simulations, the RMSE is expected to be larger for more difficult cases, and smaller for relatively easier cases.

In **Figure 2**, the horizontal axis represented the *λ* value for generating the protease *p*_3_, which meant that simulation cases from left to right had an increasing level of similarity between proteases *p*_1_ and *p*_3_, and thus had an increasing level of difficulty for deconvolving protease mixtures of the two proteases. Each curve represented a different series of simulations with a particular number of substrates. In **Figure 2**, the simulation series with a larger number of substrates led to smaller RMSEs. Note that the 2- and 3-substrates curves largely overlapped, and the 5-, 6-, and 7-substrates curves also largely overlapped. In each simulated series with a specific number of substrates, the RMSE increased in general with respect to the horizontal axis that represented an increasing level of difficulty. In the 2- and 3-substrates curves, the changes of RMSE were not monotonic. This was mainly because, with a limited number of substrates in the simulation, *λ* around 0.5 already represented quite difficult situations that led to very large RMSE with high variance. For the subsequent larger *λ* values representing even more difficult situations, the slight decrease of the subsequent RMSEs was due to the high variance when the RMSE was large. Overall, the observed RMSEs showed expected trends with respect to the level of difficulty of the simulated cases, validating that the RMSE is a useful evaluation metric.

**Figure 2.**
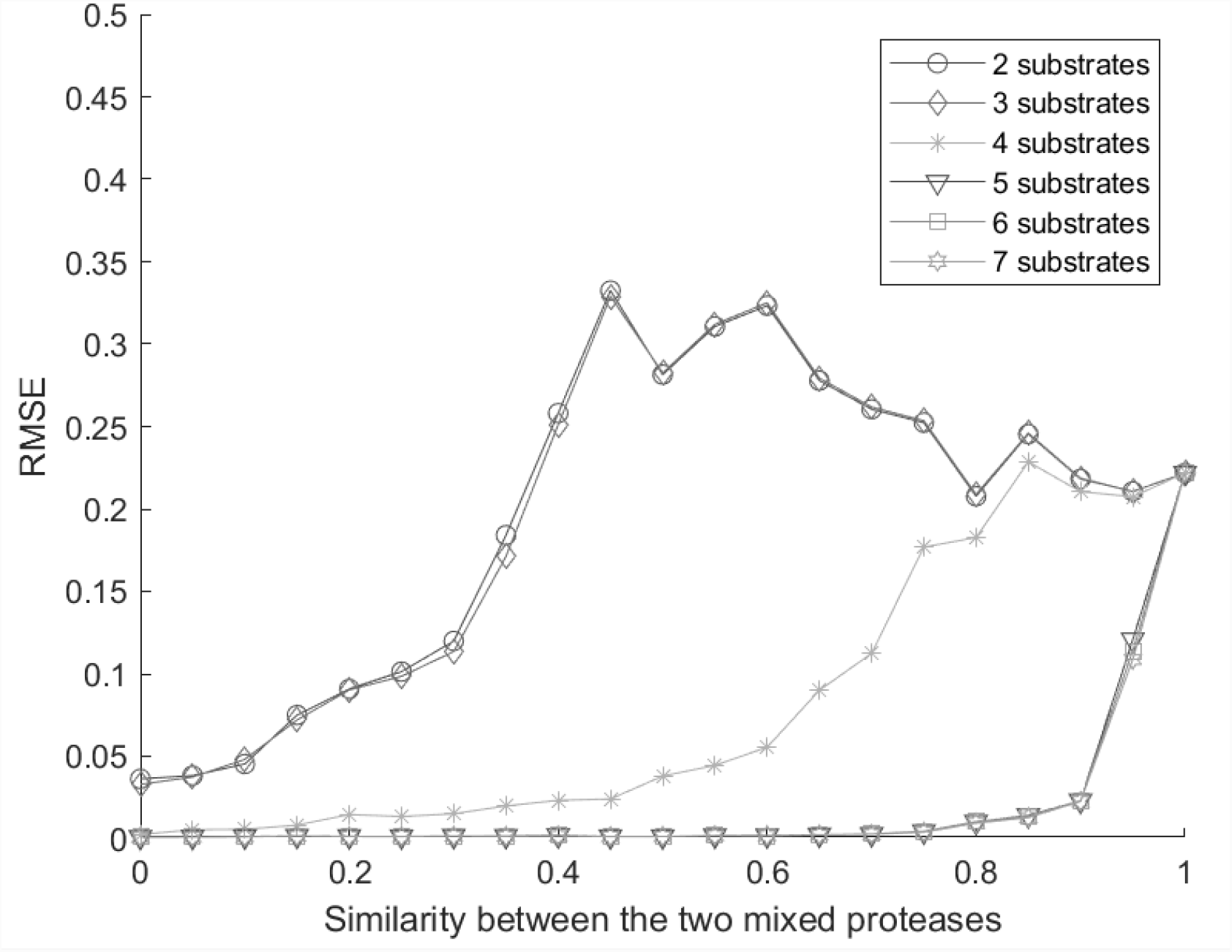
RMSE reflected the level of difficulties in deconvolution of the simulated protease mixtures. The x-axis represents the similarity between the two proteases in the mixture. A higher similarity between proteases led to a higher correlation between their cleaving dynamics, and thus higher difficulty in deconvolution. The y-axis was the RMSE of the estimated mixing coefficients. For each substrates-proteases setting, repetition P = 200 was used to calculate the corresponding reported RMSE. Each curve represented a simulation series with a different number of substrates. A smaller number of substrates corresponded to more difficult situations for deconvolution analysis.

### 3.3 Optimizing choices of substrates

To demonstrate the feasibility of optimizing choices of substrates, we considered 3 proteases and 7 substrates, with their single-protease-single-substrate kinetic parameters randomly generated. We first evaluated the accuracy for deconvolving mixtures of the 3 proteases using all 7 substrates, which led to a low RMSE shown by the right-most point on the dashed-circle line in **Figure 3**. We then removed one substrate, and evaluated the RMSE for deconvolution with 6 substrates. All 7 possibilities were evaluated, and the best RMSE was reported as the second right-most point on the dashed line, which was virtually the same as the 7-substrate scenario. We iterated this analysis, removing one substrate that had the least impact on RMSE in each iteration, until only 2 substrates remained. As shown in **Figure 3**, the RMSE remained low until the number of substrates reduced from 3 to 2. This was because the single-protease-single-substrate kinetic parameters were randomly generated, which represented 3 proteases that had independent substrate cleavage activities. In order to deconvolve 3 independent proteases, at least 3 substrates were needed.

**Figure 3.**
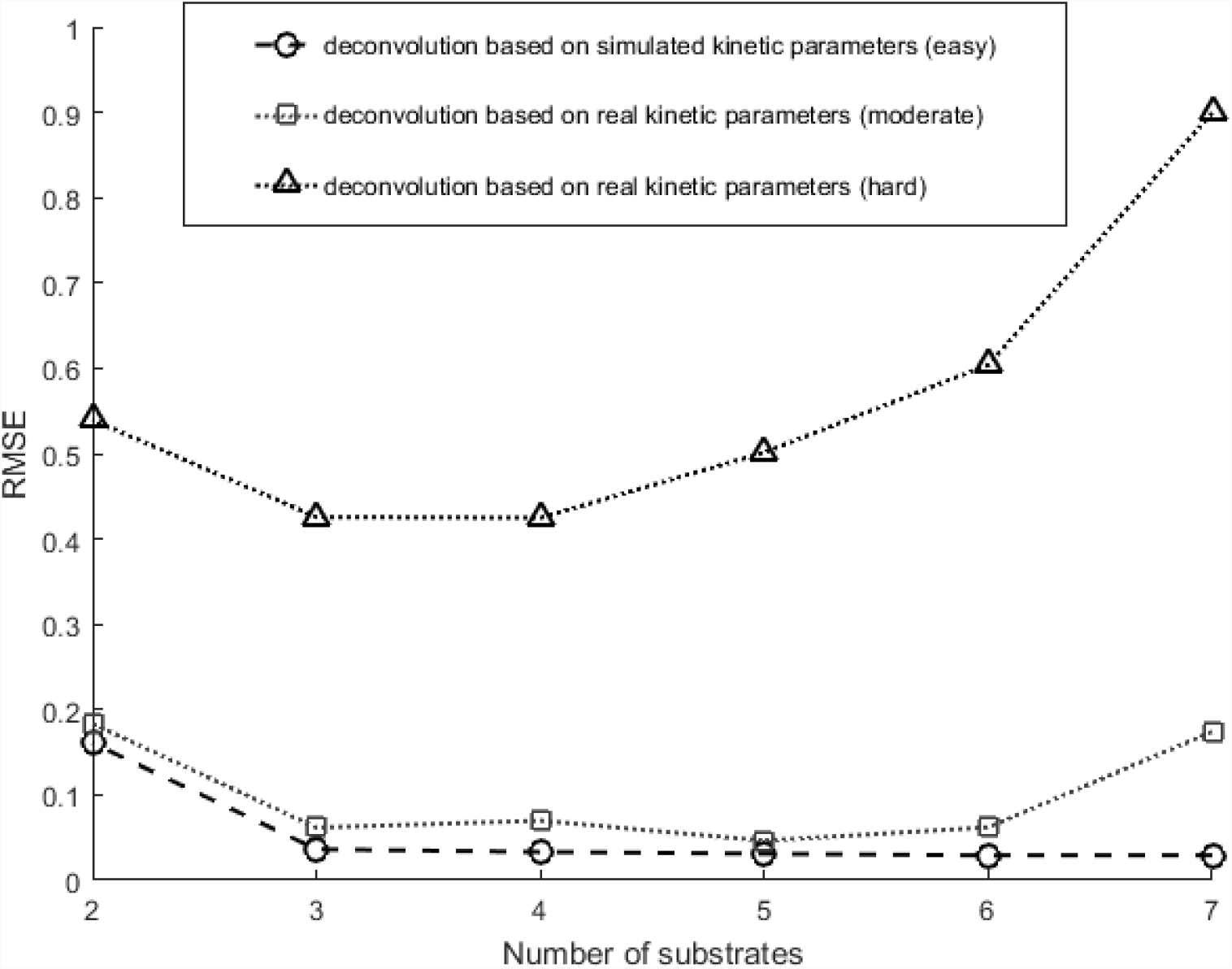
Influence of substrate library’s size on the deconvolution accuracy. The x-axis represented the number of substrates applied for deconvolution and the y-axis represented the resulting RMSE. The dashed curve with circles represented RMSE for deconvolving mixtures of three simulated proteases that are independent, and showed increased RMSE as the number of substrates decreased from 7 down to 2. The dotted curves with squares and with triangles represented RMSE for deconvolving mixtures of three real proteases that are slightly correlated and highly correlated accordingly, where the RMSEs were relatively high even when the number of substrates was 7. Repetition time P = 200 was applied for each substrates-proteases setting. Discussion regarding the decrease of RMSE when reducing the number of substrates were discussed in **Supplementary Figure S1∼S6**.

We performed two sets of similar analyses using 3 of the 7 proteases and the 7 substrates in our real experimental data in section 3.1. One set of analyses was based on proteases MASP2, C1r, and F2, which were from 3 different proteases families, and the other set of analyses was based on proteases MASP2, CFI, and CFD, which were highly correlated in terms of their substrate cleaving dynamics. The single-protease-single-substrate kinetic parameters were estimated from real experimental data. All subsequent analyses were the same as the above where kinetic parameters were randomly generated. As shown by the dotted-triangle curve in **Figure 3**, deconvolving the three highly correlated proteases was quite difficult with large the RMSE regardless of how many substrates were used. The dotted-square curve in **Figure 3** was similar to the analysis where kinetic parameters were randomly generated, indicating that the three proteases from different protease families had relative independent cleaving dynamics against the substrates. Interestingly, the performance actually improved in both dotted curves when the number of substrates reduced from 7 to 5 (or 4). This was because the first few substrates being removed had extremely similar cleaving dynamics against all the proteases (details in **Supplementary Figure S1∼S6**). Those substrates were not only uninformative but also sources of confusion for the deconvolution analysis. Therefore, effective deconvolution of protease mixtures required a decent number of substrates with uncorrelated cleaving dynamics against the proteases. However, correlated substrate cleaving dynamics is ubiquitous, especially among proteases in the same physiological family. When deconvolving mixtures containing highly correlated proteases, even a large number of substrates may not lead to satisfactory deconvolution performances. This motivated us to investigate a less ambitious goal of deconvolving protease families, instead of deconvolving individual proteases.

### 3.4 Deconvolving protease families

As mentioned above, deconvolving protease mixtures can be challenging and may require an impractically large number of substrates when proteases with highly correlated substrate cleaving dynamics exist in the mixture. Therefore, we propose to examine the possibility of clustering highly correlated proteases into families, and deconvolve the protease families by estimating the mixing concentration of the protease families.

#### 3.4.1 Deconvolving simulated protease families

We first examined a simulated scenario with 9 proteases and 7 substrates. The 9 proteases formed three families. Each family contained 3 highly correlated proteases, but the families are independent of each other. To *in silico* generate the 3 highly correlated proteases in one family, we took a similar strategy as described in Section 3.2. For each protease family, we randomly generated the kinetic parameters of three proteases (*p*_1_, *p*_2_, and *p*_3_) cleaving 7 substrates, and then generated two proteases *p*_4_ and *p*_5_ that correlated *p*_1_ using the following combinations *p*_4_ = *λp*_1_ + (1 − *λ*)*p*_2_, and *p*_5_ =*λp*_1_ + (1 − *λ*)*p*_3_. Proteases *p*_1_, *p*_4_ and *p*_5_ form the family. Here, *λ* was either 0.9 or 0.6, representing a proteases family containing highly correlated proteases or moderately correlated proteases. We repeated the above three times to generate the kinetic parameters for the three families of proteases.

After generating the single-protease-single-substrate kinetic parameters with family structures, we simulated data for the single-protease-single-substrate setting and the multi-proteases-multi-substrates setting. We then evaluated the performance for deconvolving the 9 individual proteases using 3, 5, or 7 substrates. **Figure 4a and 4c** showed scatter plots of the true simulated mixing coefficients versus the estimated mixing coefficients, where the estimation performance was poor. The only exception was the case in the third plot of **Figure 4c**, where the protease was moderately correlated (*λ* = 0.6) and the number of substrates was 7. This was the least challenging case simulated here for deconvolving individual proteases, where the estimated mixing coefficients roughly tracked the true mixing coefficients. Overall, with the presence of correlated proteases, although the protease families were independent, it was difficult to deconvolve the mixing coefficients of the individual proteases.

**Figure 4.**
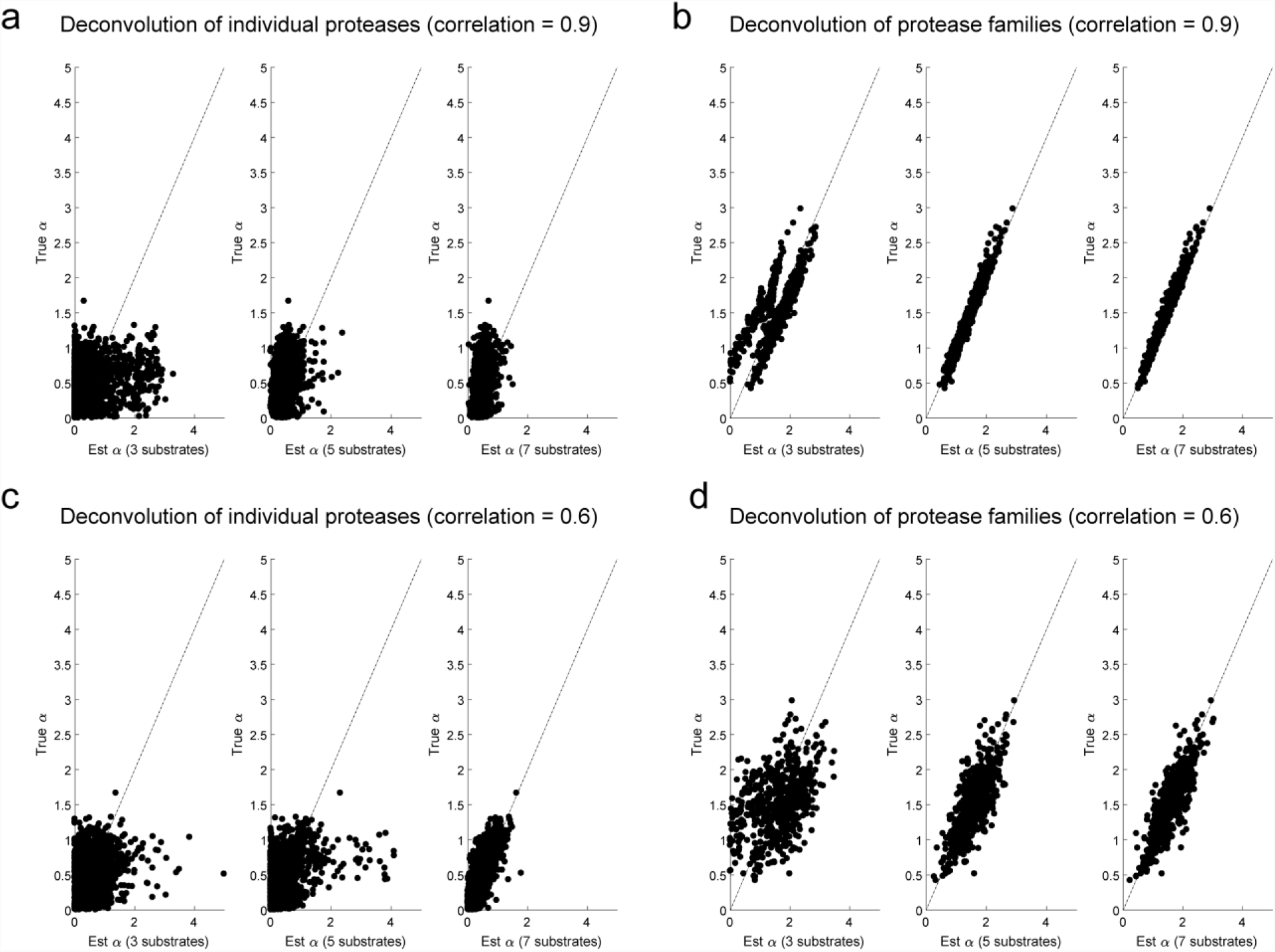
Comparison between deconvolution of individual proteases and deconvolution of protease families. The x axis represented the estimated mixing coefficients. The y axis represented the true simulated mixing coefficients. Consider mixtures of 9 proteases from 3 independent families that contain highly correlated proteases within each family. **(a)** When deconvolving the 9 individual proteases, the estimated mixing coefficients for the individual proteases showed poor agreements with their true simulated values, regardless of whether 3, 5, or 7 substrates were used. **(b)** When deconvolving the 3 protease families, the estimated mixing coefficients for the protease families showed high agreement with the sum of the family members’ true mixing coefficients. **(c-d)** Simulation of protease families that contain moderately correlated proteases showed similar results, where deconvolution of individual proteases was difficult but deconvolution of protease families was accurate.

Using the same simulated data as above, we evaluated the possibility for deconvolving protease families. In order to perform deconvolution at the protease family level, we used Equation (4) to estimate one set of kinetic parameters for each family, by treating the simulated single-protease-single-substrate data for protease members in the same family as replicates of a “representative” protease for the family. After estimating the kinetic parameters for the three protease families, we then optimized Equation (5) to estimate the mixing coefficient of the protease families. Ideally, the estimated mixing coefficient for one protease family should approximate the sum of the true mixing coefficients of members in the family, which was indeed what we observed in the results shown in **Figure 4b and 4d**. Meanwhile, the performance difference between **Figure 4b and 4d** indicated that successful deconvolution of protease families required members within each family to be decently correlated. This analysis demonstrated the feasibility of accurately deconvolving protease families, while the deconvolution of individual proteases was difficult.

#### 3.4.2 Deconvolving protease families derived from real data

To further validate the idea of deconvolving protease families, we used kinetic parameters estimated from the real single-protease-single-substrate data in Section 3.1. We performed hierarchical clustering using the single-protease-single-substrate cleaving data and decided to group the 7 individual proteases 4 families (as shown in **Figure 5**). After that, we performed the same analysis as above, including simulation of the multi-protease-multi-substrate data, estimation of the kinetic parameters for the protease families, and estimation of the mixing coefficients for the protease families. We also performed deconvolution for the individual proteases. As shown in **Figure 6**, the deconvolution of the individual proteases performed poorly, while the estimated mixing coefficients of the protease families decently tracked the simulated true mixing coefficients. The deconvolution accuracy of these protease families was lower than the above analysis of randomly generated kinetic parameters. This was because the randomly generated kinetic parameters led to simulated protease families that were independent of each other, whereas the protease families derived from real data were not as independent. Since human proteases are known to organize in a nested hierarchy of protease families and subfamilies, the correlation among them makes deconvolution of individual protease cumbersome and impractical. This analysis provides a practical alternative for sensing and characterizing protease activities at the level of protease families.

**Figure 5.**
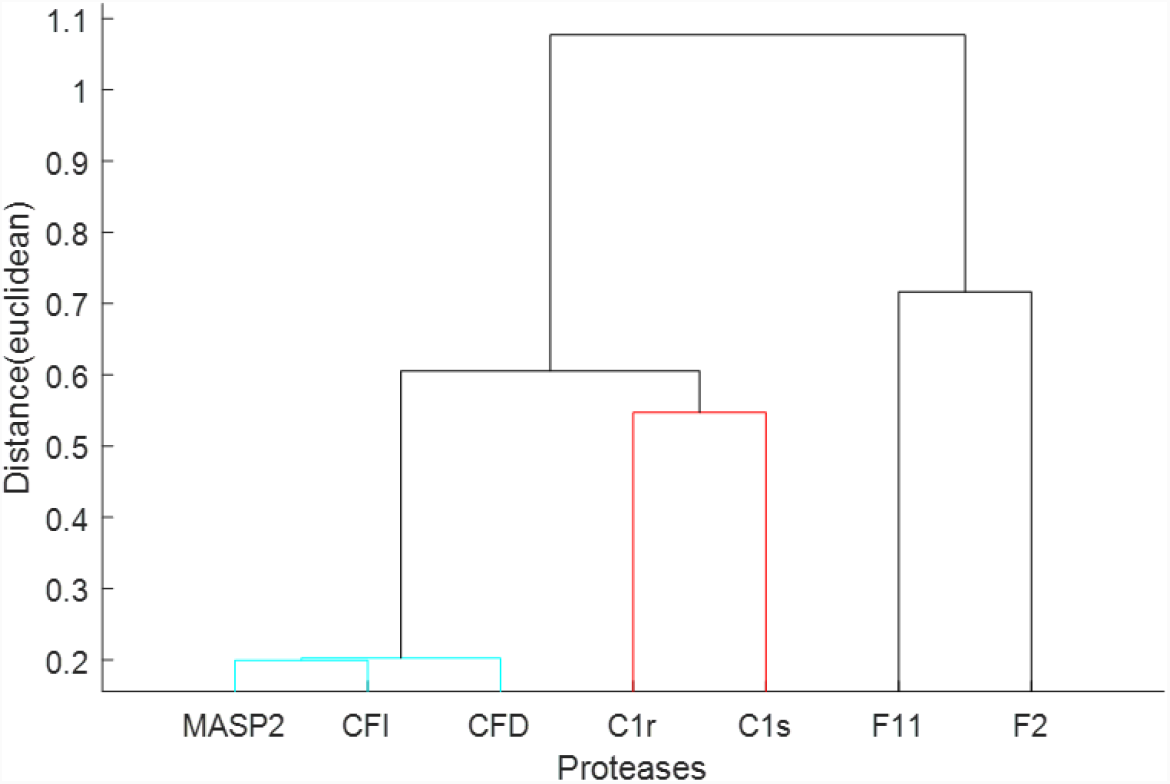
Hierarchical clustering of individual proteases led to 4 families. MASP2+CFI+CFD formed the most strongly correlated family, C1r + C1s formed the moderately correlated family, and F11 and F2 each served as its own family.

**Figure 6.**
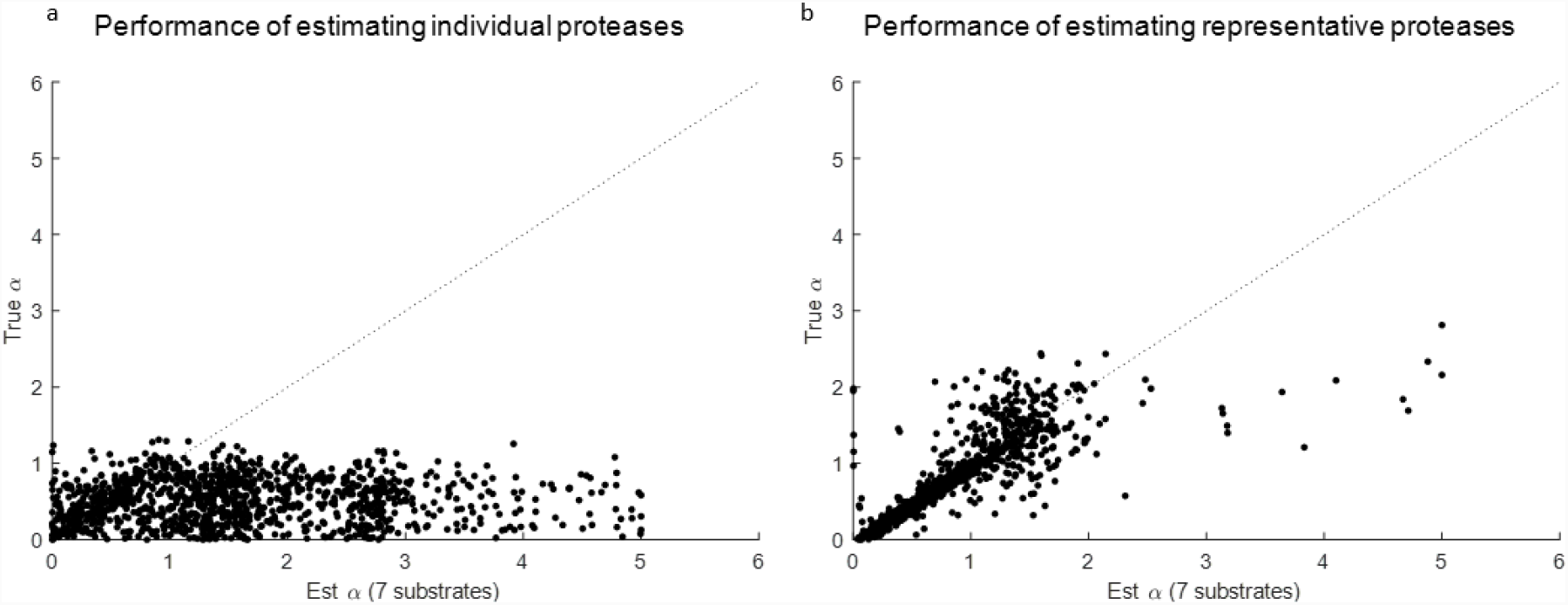
Performances of estimating representative families and individual proteases using the same 7 proteases. **(a)** Deconvolution of 7 individual proteases was no better than a random guess since the scatter dots spread out widely on the horizontal direction. **(b)** After clustering 7 individual proteases into 4 protease families as shown in Figure 5, deconvolution accuracy increased, with mixing coefficients of protease families being close to the sum of individual mixing coefficients within according protease families.

## 4 Conclusion

While significant advances were made by using genome-wide sequencing techniques, the lack of correlation between expression and activity is well-known (*14*). Further, the primary drivers of all physiological processes in health and disease are enzymes (e.g., proteases, kinases), meaning the most valuable information is stored in the real-time activity of these proteins. One of the main challenges in scaling up to multiplexed libraries for protease activity analysis is substrate design, due to the difficulty of screening for specific substrates. Due to the promiscuity of proteases, the goal of designing substrates with both high sensitivity and high specificity for all human proteases is experimentally extremely challenging, if not infeasible. We overcame this challenge by using a set of nonspecific substrates, which required minimal design effort, and instead leveraged computational deconvolution. Here, we present a framework for evaluating the activity contributions of individual proteases within a complex mixture.

Fundamentally, however, human proteases are organized in a hierarchy of protease families that consist of proteases with highly correlated activities against many substrates. Deconvolution of highly correlated protease would require an impractically large number of substrates. Therefore, we propose to cluster highly correlated proteases into families, and perform deconvolution of the protease families. We demonstrate the feasibility of accurately deconvolving protease families when the deconvolution of individual proteases was difficult. In practice, this allows for rapid characterization and investigation of physiologically relevant protease families, which can effectively screen the entire protease landscape before homing in on specific protease targets.

## Supplementary

**Figure S1.**
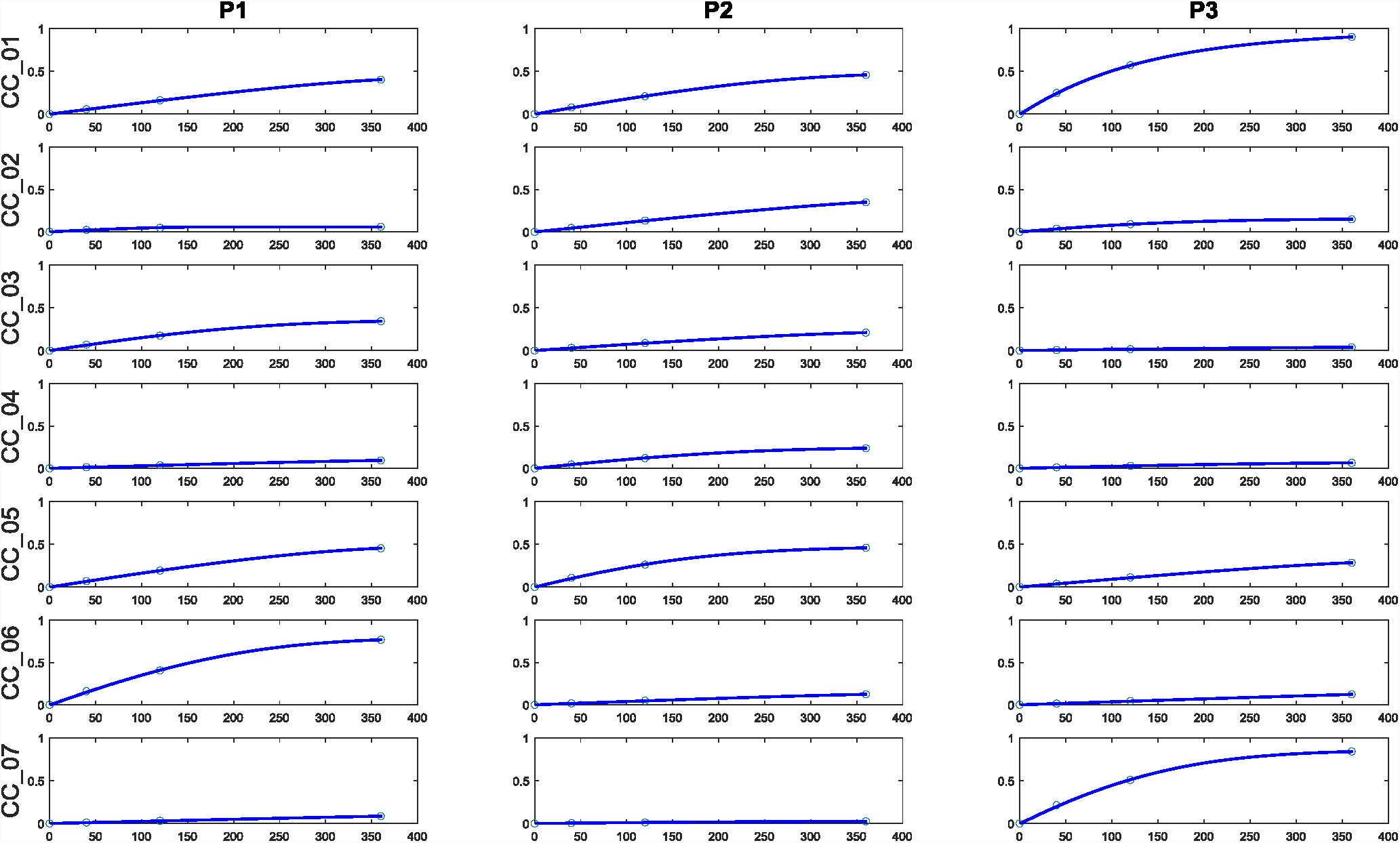
Substrate cleavage dynamics generated by simulated kinetics parameters for a 7-substrates-3-proteases setting. The x-axis represented the reaction time, and the y-axis represented the amount of reaction product (cleaved substrates). The amounts of cleaved substrates at t = [0 30 60 300] were collected for subsequent analysis. The 3 generated proteases showed independence in terms of their substrate cleaving dynamics.

**Figure S2.**
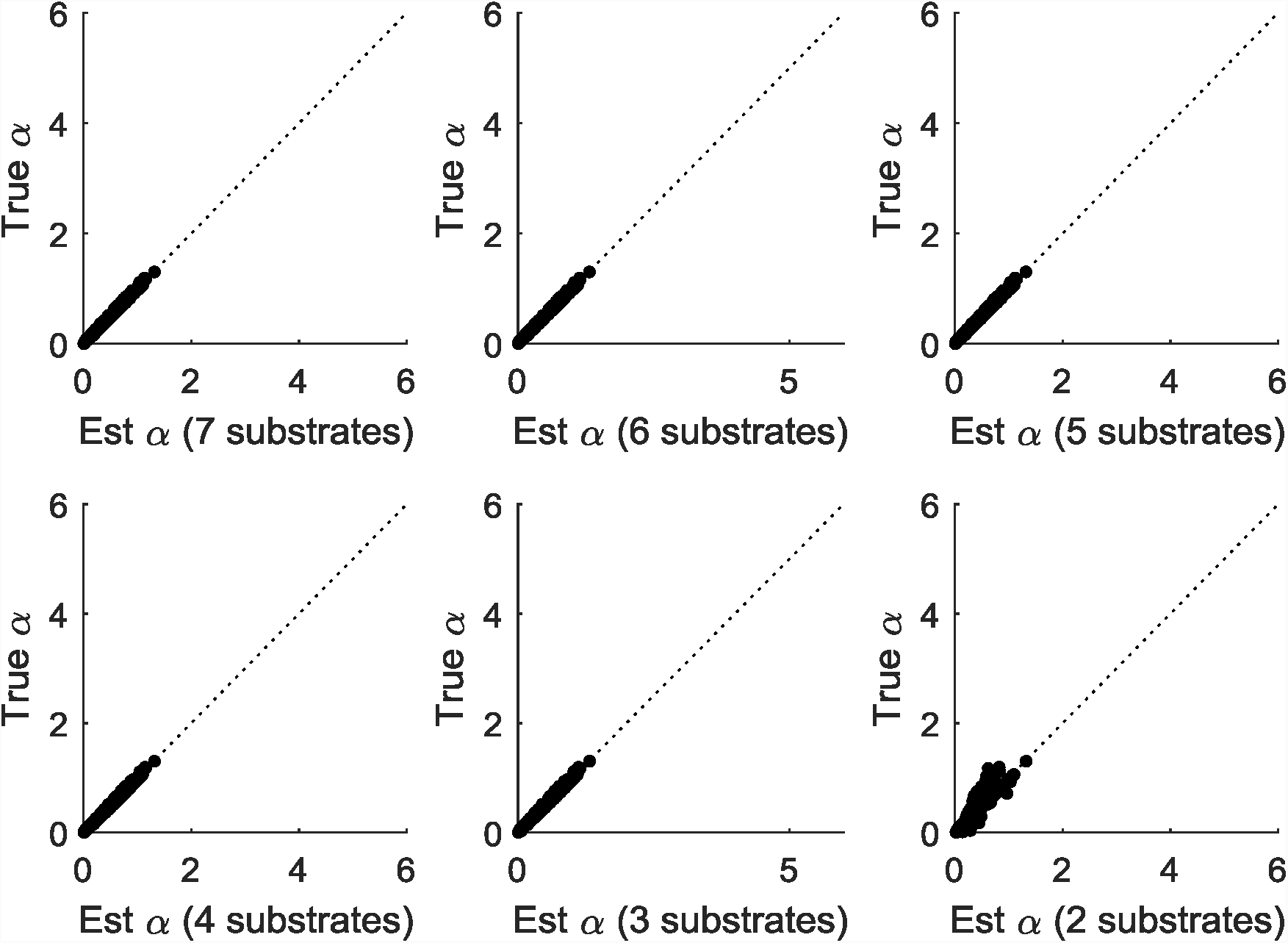
Deconvolution performance of the 7-substrates-3-proteases setting using simulated kinetic parameters. The x-axis represented the estimated mixing coefficients. The y-axis represented the true simulated mixing coefficients. The deconvolution performance stayed quite decent along the way of reducing the number of substrates from 7 to 3, and showed a decrease in accuracy when reducing from 3 substrates to 2 substrates. To deconvolute 3 independent proteases, the sufficient number of proteases was 3. The deconvolution difficulty level of this independent-proteases setting was regarded as “easy” in Figure 3.

**Figure S3.**
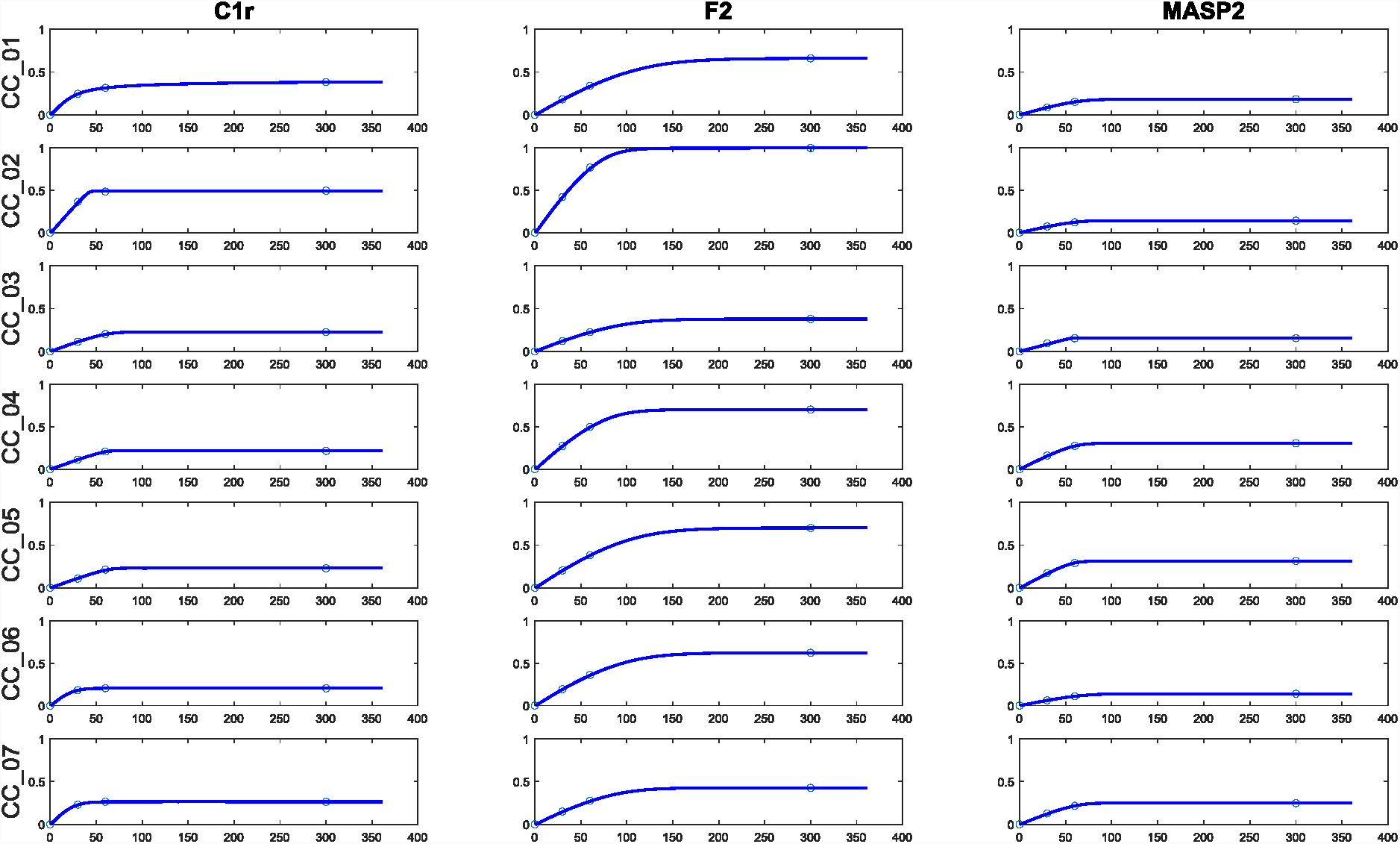
Substrate cleavage dynamics generated by estimated kinetics parameters from a 7-substrates-3-proteases setting in the real experimental data. The involved proteases (from different families) were C1r, F2, and MASP2. The x-axis represented the reaction time, and the y-axis represented the amount of reaction product (cleaved substrates). The amounts of cleaved substrates at t = [0 30 60 300] were collected for subsequent analysis. The three proteases showed a moderate level of correlation in terms of their substrate cleaving dynamics, especially for substrates CC_03 and CC_07.

**Figure S4.**
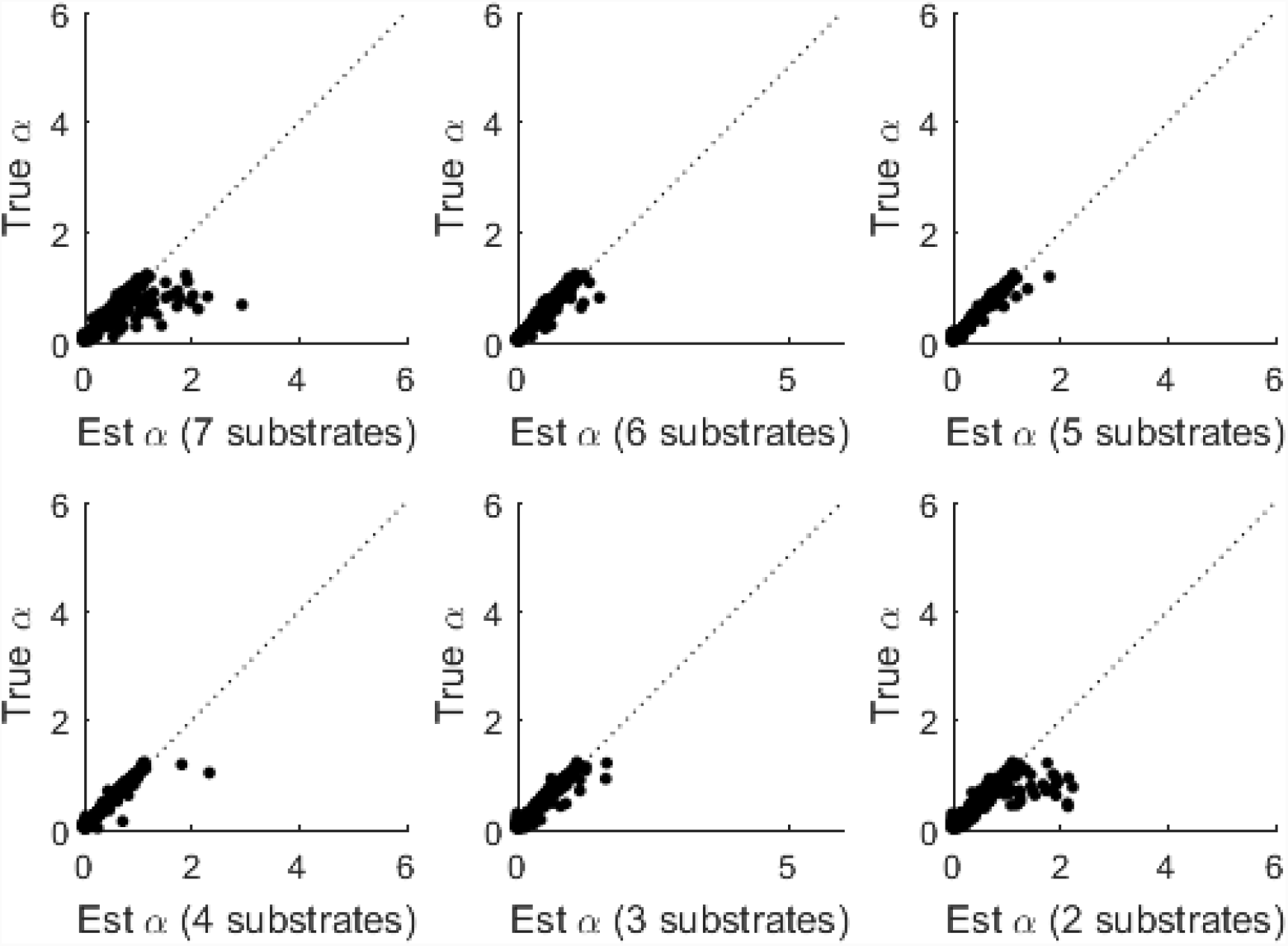
Deconvolution performance of the 7-substrates-3-proteases setting using kinetic parameters estimated from real experimental data. The involved proteases were C1r, F2, and MASP2. The x-axis represented the estimated mixing coefficients. The y-axis represented the true simulated mixing coefficients. The deconvolution accuracy increased when reducing from 7 substrates to 6 and to 5 substrates, and decreased when reducing from 4 substrates to 3 and 2 to substrates. This counterintuitive observation could be explained by the high correlation among three proteases when cleaving substrates CC_03 and CC_07 in Figure S3. Removing CC_03 and CC_07, which could be regarded as “noise”, decreased the overall level of correlation and thus increased the deconvolution performance (from 7 substrates to 6 and to 5 substrates). With the correlation among proteases being reduced, further removing the number of substrates from 4 to 3 and to 2 decreased the deconvolution accuracy due to loss of information. The deconvolution difficulty level of this moderately-correlated setting was regarded as “moderate” in Figure 3.

**Figure S5.**
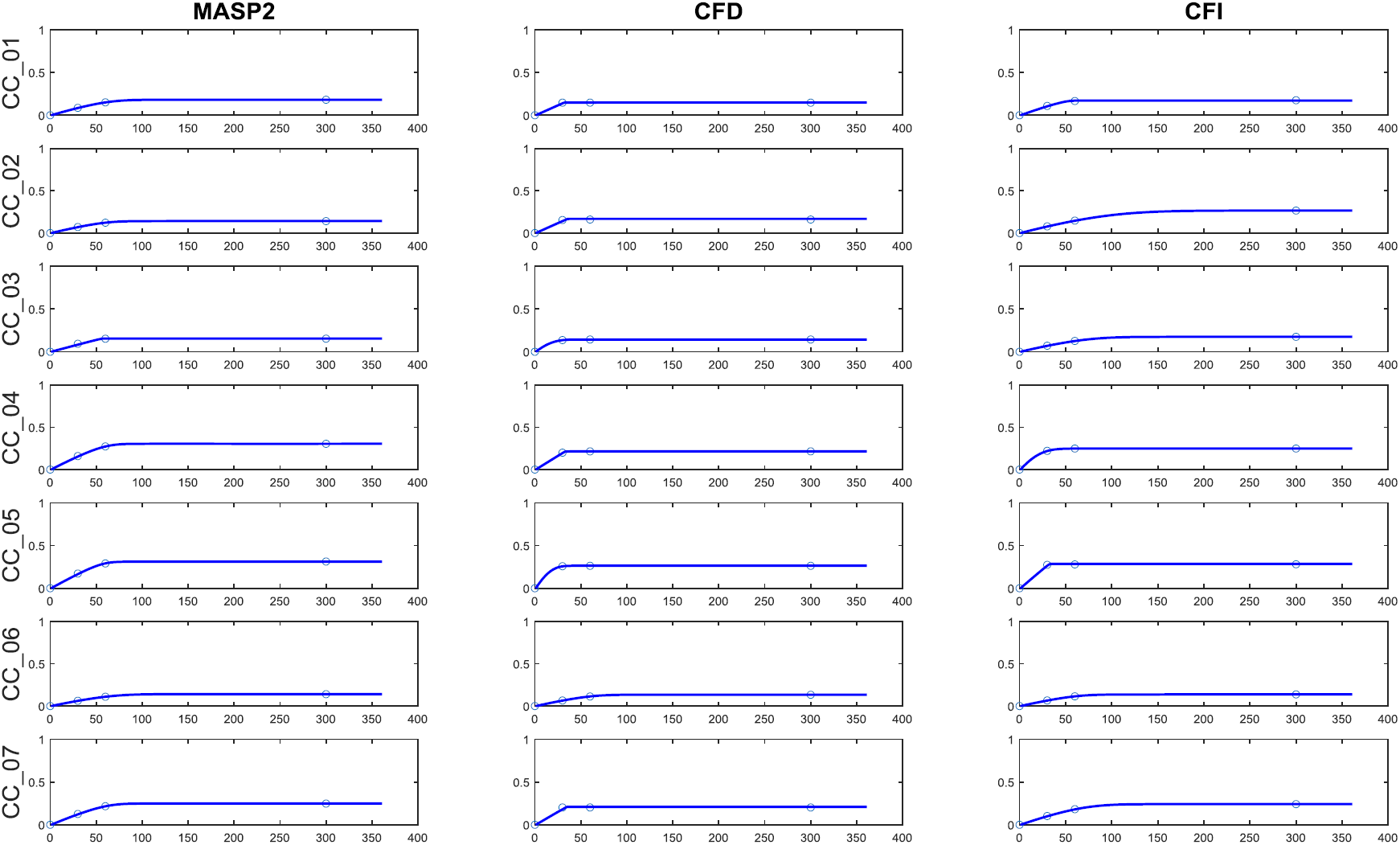
Substrate cleavage dynamics generated by estimated kinetics parameters from a 7-substrates-3-proteases setting in the real experimental data. The involved proteases (from the same family) were MASP2, CFD, and CFI. The x-axis represented the reaction time, and the y-axis represented the amount of reaction product (cleaved substrates). The amounts of cleaved substrates at t = [0 30 60 300] were collected for subsequent analysis. The three proteases showed a high level of correlation in terms of their substrate cleaving dynamics, for almost all substrates.

**Figure S6.**
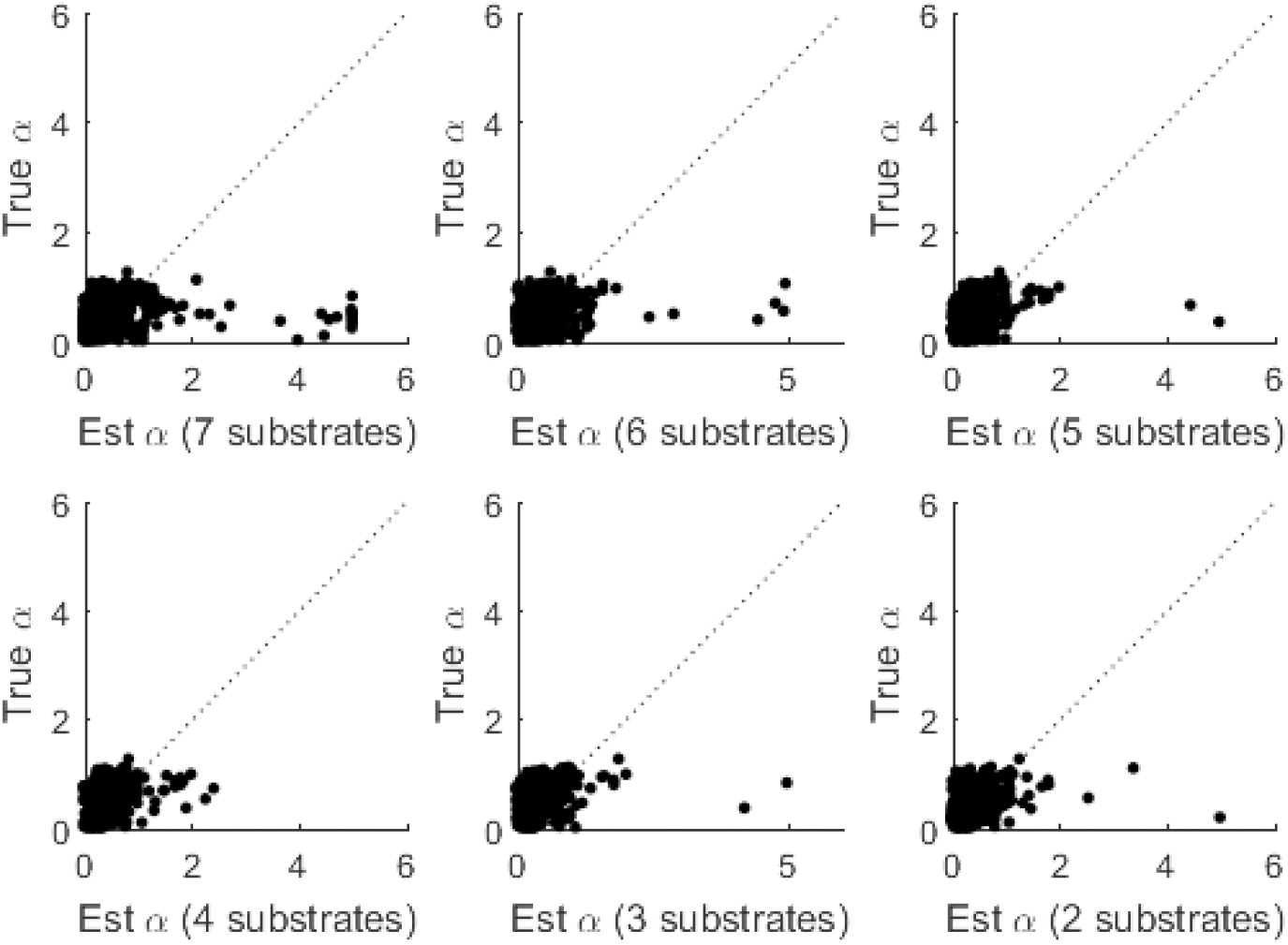
Deconvolution performance of the 7-substrates-3-proteases setting using kinetic parameters estimated from real experimental data. The involved proteases (from the same family) were MASP2, CFD, and CFI. The x-axis represented the estimated mixing coefficients. The y-axis represented the true simulated mixing coefficients. The deconvolution stayed poor regardless of the number of substrates being applied. Similar to Figure S4, the slight increase in deconvolution accuracy when reducing from 7 substrates down to 4 substrates was observed again. The deconvolution difficulty level of this highly-correlated setting was regarded as “hard” in Figure 3.

## References

1. P. I. Bird, J. A. Trapani, J. A. Villadangos, Endolysosomal proteases and their inhibitors in immunity. Nature Reviews Immunology 9, 871 (2009).

2. G. A. de Souza, L. M. F. de Godoy, M. Mann, Identification of 491 proteins in the tear fluid proteome reveals a large number of proteases and protease inhibitors. Genome Biology 7, R72 (2006).

3. D. Frees, L. Brondsted, H. Ingmer, Bacterial proteases and virulence. Sub-cellular biochemistry 66, 161–192 (2013).

4. K. M. Heutinck, I. J. ten Berge, C. E. Hack, J. Hamann, A. T. Rowshani, Serine proteases of the human immune system in health and disease. Molecular immunology 47, 1943–1955 (2010).

5. Y. Hua, S. Nair, Proteases in cardiometabolic diseases: Pathophysiology, molecular mechanisms and clinical applications. Biochimica et biophysica acta 1852, 195–208 (2015).

6. H. Ingmer, L. Brondsted, Proteases in bacterial pathogenesis. Research in microbiology 160, 704–710 (2009).

7. J. E. Koblinski, M. Ahram, B. F. Sloane, Unraveling the role of proteases in cancer. Clinica chimica acta; international journal of clinical chemistry 291, 113–135 (2000).

8. C. Lopez-Otin, J. S. Bond, Proteases: multifunctional enzymes in life and disease. The Journal of biological chemistry 283, 30433–30437 (2008).

9. J. G. Pérez-Silva, Y. Español, G. Velasco, V. Quesada, The Degradome database: expanding roles of mammalian proteases in life and disease. Nucleic Acids Research 44, D351–D355 (2016).

10. L. E. Sanman, M. Bogyo, Activity-based profiling of proteases. Annual review of biochemistry 83, 249–273 (2014).

11. C. M. Overall, C. P. Blobel, In search of partners: linking extracellular proteases to substrates. Nature Reviews Molecular Cell Biology 8, 245 (2007).

12. S. Miyamoto et al., Discrepancies between the gene expression, protein expression, and enzymatic activity of thymidylate synthase and dihydropyrimidine dehydrogenase in human gastrointestinal cancers and adjacent normal mucosa. International journal of oncology 18, 705–713 (2001).

13. O. Takumi et al., [Correlation between enzymatic activity and gene expression of orotate phosphoribosyl transferase (OPRT) in colorectal cancer]. Gan to kagaku ryoho. Cancer & chemotherapy 29, 2515–2519 (2002).

14. J. Yin et al., Study on the Correlation between Gene Expression and Enzyme Activity of Seven Key Enzymes and Ginsenoside Content in Ginseng in Over Time in Ji’an, China. International Journal of Molecular Sciences 18, 2682 (2017).

15. J. S. Dudani, P. K. Jain, G. A. Kwong, K. R. Stevens, S. N. Bhatia, Photoactivated Spatiotemporally-Responsive Nanosensors of in Vivo Protease Activity. ACS Nano 9, 11708–11717 (2015).

16. L. E. Edgington, M. Verdoes, M. Bogyo, Functional imaging of proteases: recent advances in the design and application of substrate-based and activity-based probes. Current opinion in chemical biology 15, 798–805 (2011).

17. M. Fonovic, M. Bogyo, Activity based probes for proteases: applications to biomarker discovery, molecular imaging and drug screening. Current pharmaceutical design 13, 253–261 (2007).

18. B. A. Holt et al., Fc microparticles can modulate the physical extent and magnitude of complement activity. Biomaterials science 5, 463–474 (2017).

19. B. A. Holt, Q. D. Mac, G. A. Kwong, Nanosensors to Detect Protease Activity In Vivo for Noninvasive Diagnostics. JoVE, e57937 (2018).

20. E. J. Kwon, J. S. Dudani, S. N. Bhatia, Ultrasensitive tumour-penetrating nanosensors of protease activity. Nature biomedical engineering 1, (2017).

21. G. A. Kwong et al., Mathematical framework for activity-based cancer biomarkers. Proceedings of the National Academy of Sciences 112, 12627–12632 (2015).

22. D. K. Nomura, M. M. Dix, B. F. Cravatt, Activity-based protein profiling for biochemical pathway discovery in cancer. Nature reviews. Cancer 10, 630–638 (2010).

23. S. Schuerle, J. S. Dudani, M. G. Christiansen, P. Anikeeva, S. N. Bhatia, Magnetically Actuated Protease Sensors for in Vivo Tumor Profiling. Nano Letters 16, 6303–6310 (2016).

24. V. Stein, K. Alexandrov, Protease-based synthetic sensing and signal amplification. Proceedings of the National Academy of Sciences 111, 15934–15939 (2014).

25. T.-L. To et al., Rationally designed fluorogenic protease reporter visualizes spatiotemporal dynamics of apoptosis in vivo. Proceedings of the National Academy of Sciences 112, 3338–3343 (2015).

26. J. Villanueva, A. Nazarian, K. Lawlor, P. Tempst, Monitoring Peptidase Activities in Complex Proteomes by MALDI-TOF Mass Spectrometry. Nature protocols 4, 1167–1183 (2009).

27. J. Villanueva et al., Differential exoprotease activities confer tumor-specific serum peptidome patterns. The Journal of Clinical Investigation 116, 271–284 (2006).

28. A. D. Warren et al., Disease detection by ultrasensitive quantification of microdosed synthetic urinary biomarkers. Journal of the American Chemical Society 136, 13709–13714 (2014).

29. Q. D. Mac et al., Non-invasive early detection of acute transplant rejection via nanosensors of granzyme B activity. Nature biomedical engineering, (2019).

30. B. A. Holt, G. A. Kwong, Bacterial defiance as a form of prodrug failure. bioRxiv, 556951 (2019).

31. G. A. Kwong et al., Mass-encoded synthetic biomarkers for multiplexed urinary monitoring of disease. Nat Biotech 31, 63–70 (2013).

32. M. A. Miller et al., Proteolytic Activity Matrix Analysis (PrAMA) for simultaneous determination of multiple protease activities. Integr Biol (Camb) 3, 422–438 (2011).

33. L. Menten, M. Michaelis, Die kinetik der invertinwirkung. Biochem Z 49, 333–369 (1913).

34. J. P. Keener, J. Sneyd, Mathematical physiology. (Springer, 1998), vol. 1.

35. S.-P. Han, A globally convergent method for nonlinear programming. Journal of optimization theory and applications 22, 297–309 (1977).

36. M. J. Powell, in Numerical analysis. (Springer, 1978), pp. 144–157.

37. M. J. Powell, in Nonlinear programming 3. (Elsevier, 1978), pp. 27–63.

38. T. Hastie, R. Tibshirani, J. Friedman, The Elements of Statistical Learning. (2009).

